# True S-cones are concentrated in the ventral mouse retina for color detection in the upper visual field

**DOI:** 10.1101/2020.03.20.999979

**Authors:** Francisco M. Nadal-Nicolás, Vincent P. Kunze, John M. Ball, Brian Peng, Akshay Krisnan, Gaohui Zhou, Lijin Dong, Wei Li

## Abstract

Color, an important visual cue for survival, is encoded by comparing signals from photoreceptors with different spectral sensitivities. The mouse retina expresses a short wavelength-sensitive and a middle/long wavelength-sensitive opsin (S- and M-opsin), forming opposing, overlapping gradients along the dorsal-ventral axis. Here, we analyzed the distribution of all cone types across the entire retina for two commonly used mouse strains. We found, unexpectedly, that ‘true’ S-cones (S-opsin only) are highly concentrated (up to 30% of cones) in ventral retina. Moreover, S-cone bipolar cells (SCBCs) are also skewed towards ventral retina, with wiring patterns matching the distribution of true S-cones. In addition, true S-cones in the ventral retina form clusters, which may augment synaptic input to SCBCs. Such a unique true S-cone pattern forms a basis for mouse color vision, likely reflecting evolutionary adaption to enhance color coding for the upper visual field suitable for mice’s habitat and behavior.

## 1. INTRODUCTION

Topographic representation of the visual world in the brain originates from the light-sensitive photoreceptors in the retina (Rhim et al., 2017). Although the neuronal architecture of the retina is similar among different vertebrates, the numbers and distributions of photoreceptors vary considerably (Hunt and Peichl, 2014). Such patterns have been evolutionarily selected, adapting to the animal’s unique behavior (diurnal or nocturnal) and lifestyle (prey or predator) for better use of the visual information in the natural environment (Dominy and Lucas, 2001; Gerl and Morris, 2008; Peichl, 2005). Color, an important visual cue for survival, is encoded by comparing signals carried by photoreceptors with different spectral preferences (Baden and Osorio, 2019). While trichromatic color vision is privileged for some primates, dichromatic vision is the evolutionarily ancient retinal circuit common to most mammals (Marshak and Mills, 2014; Puller and Haverkamp, 2011; Jacobs, 1993). The mouse retina, a model widely used for vision research, expresses two types of opsins, S- and M-opsin, that peak at 360 nm and 508 nm respectively (Jacobs et al., 1991; Nikonov et al., 2006). The expression patterns of these two opsins form opposing and overlapping gradients along the dorsal-ventral axis, resulting in a majority of cones expressing both opsins (herein either “mixed cones” or M^+^S^+^) (Applebury et al., 2000; Ng et al., 2001; Wang et al., 2011). While co-expression of both opsins broadens the spectral range of individual cones and improves perception under varying conditions of ambient light (Chang et al., 2013), which provides for a nearly optimal achromatic contrast above and below the horizon (Baden et al., 2013), it does pose a challenge for color-coding, particularly so for mixed cones that lack narrow spectral tuning. However, it has been discovered that a small population of cones only expressing S-opsin (“true S-cones”, or S^+^M^-^). These true S-cones are thought to be evenly distributed across the retina and to be critical for encoding color, especially in the dorsal retina where they are quasi-evenly distributed in a sea of cones expressing only M-opsin (“true M-cones”, or M^+^S^-^), a pattern akin to mammalian retinas in general (Haverkamp et al., 2005; Wang et al., 2011). Nonetheless, subsequent physiological studies revealed that color-opponent retinal ganglion cells (RGCs) are more abundant in the dorsal-ventral transition zone (Warwick et al., 2018) and the ventral retina (Joesch and Meister, 2016). Intriguingly, a behavior-based mouse study demonstrated that their ability to distinguish color is restricted to the ventral retina (Denman et al., 2018). These results prompt us to study, at the single-cell level and across the whole retina, the spatial distributions of cone types with different opsin expression configurations in order to better understand the anatomical base for the unique color-coding scheme of the mouse retina.

## 2. RESULTS AND DISCUSSION

### 2.1. True S-cones are highly concentrated in the ventral retina of pigmented mouse

In mouse retina, the gradients of S- and M-opsin expression along the dorsal-ventral axis have been well documented (Fig. 1A-B), but the distribution of individual cone types with different combinations of opsin expression across the whole retina has not been characterized (but see Baden et al., 2013, which we discuss below). We developed a highly reliable algorithm (Fig. S1A-B) to automatically quantify different cone types (true S; true M, and mixed cones, Fig. S1C) based on high-resolution images of entire flat-mount retinas immunolabeled with S- and M-opsin antibodies. Surprisingly, instead of finding an even distribution of true S-cones as previously believed (Baden et al., 2013; Haverkamp et al., 2005; Wang et al., 2011), we found the ventral region had much more numerous S-cones (∼30% of the local cone population; Fig. 1C left, Table S1A) than did the dorsal region (∼1%). This result is evident from density plots of cone types, showing highly concentrated true S-cones in the ventral retina (Fig. 2A, left column, bottom row).

**Figure 1.**
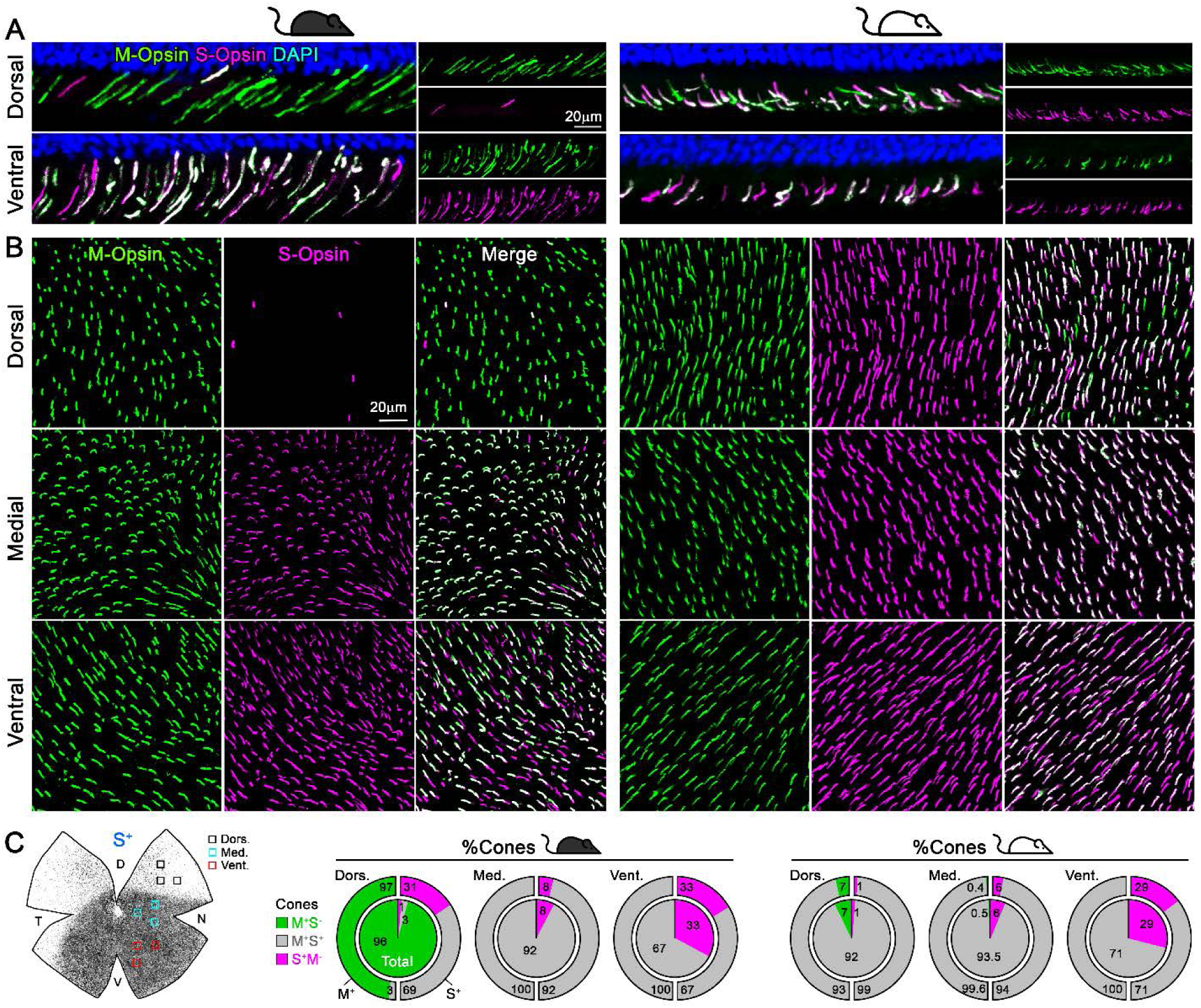
Cone outer segments across retinal areas. Immunodetection of M and S wavelength-sensitive opsins in retinal sections (A) and flat-mount retinas (B) in two mouse strains (pigmented and albino mice, left and right columns respectively). (C) Retinal scheme of S-opsin expression used for image sampling to quantify and classify cones in three different retinal regions. Pie graphs showing the percentage of cones manually classified as M^+^S^-^ (true M, green), S^+^M^-^ (true S, magenta) and M^+^S^+^ (mixed, gray) based on the opsin expression in different retinal areas. The outer rings show the relative proportions of M-opsin^+^ or S-opsin^+^ cones to mixed (gray) or true M- or S-cones (green or magenta respectively). Black mouse: pigmented mouse strain (C57BL6), white mouse: albino mouse strain (CD1).

**Figure 2.**
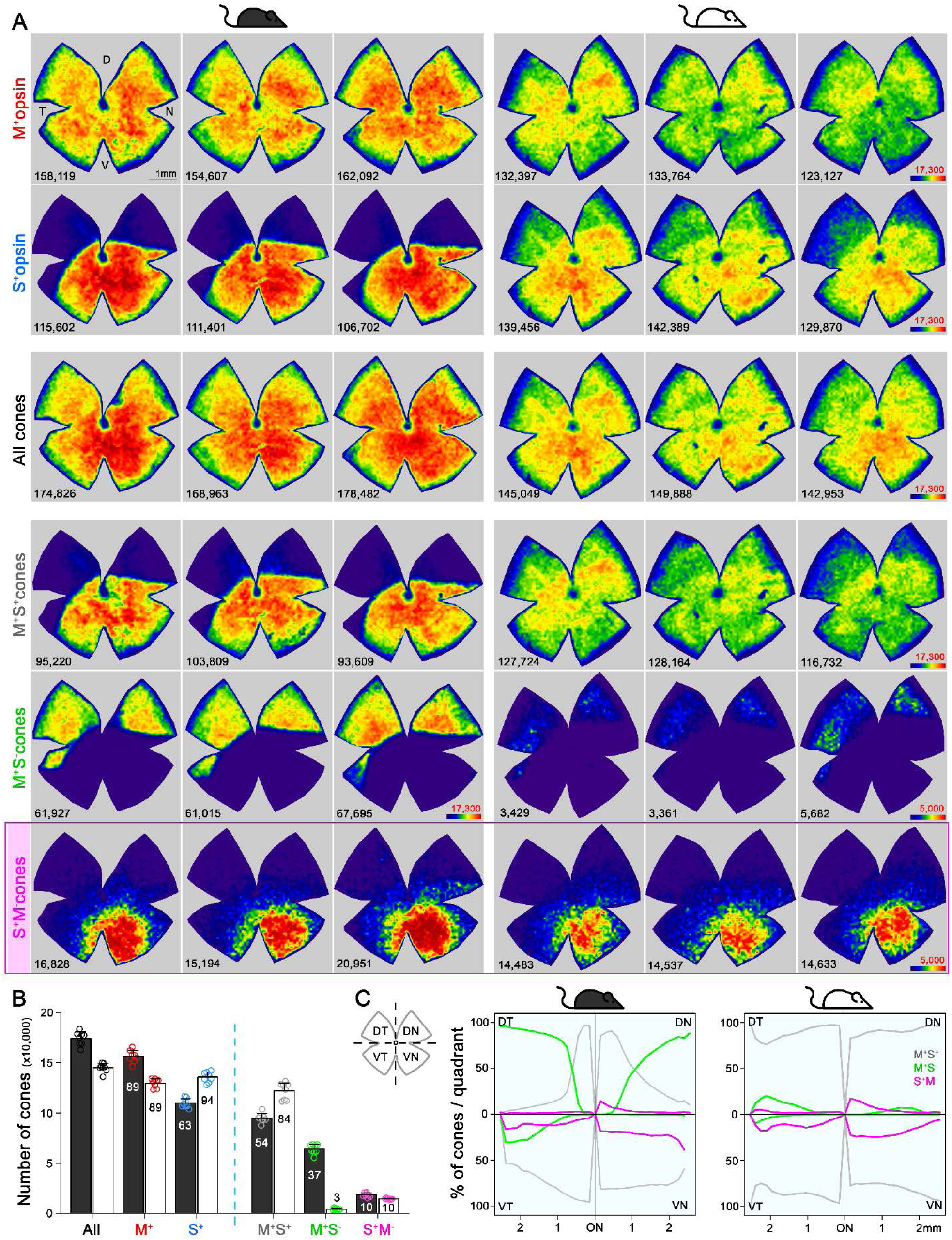
Topography and total number of different opsins (M^+^, S^+^) and cone-type populations in the whole mouse retina. (A) Density maps depicting the distributions of different cone populations classified anatomically as: All, M^+^S^+^ (mixed) M^+^S^-^ (true M), S^+^M^-^ (true S) in pigmented and albino mice (left and right side respectively). Each column shows different cone populations from the same retina and, at the bottom of each map is shown the number of quantified cones. Color scales are shown in the right panel of each row (from 0 [purple] to 17,300 or 5,000 cones/mm^2^ [dark red] for all cone types except to the true S-cones and true M-cone in the albino strain). Retinal orientation depicted by D: dorsal, N: nasal, T: temporal, V: ventral. (B) Histogram showing the mean ± standard deviation of different cone subtypes (Table 1B). The percentages of each cone subtype are indicated inside of each bar, where 100% indicates the total of the ‘all cones’ group. (C) Opsin expression profile across the different retinal quadrants (retinal scheme, DT: dorsotemporal, DN: dorsonasal, VT: ventrotemporal, VN: ventronasal). Line graphs showing the spatial profile of relative opsins expression (mixed [gray], true M-[green], true S-cones [magenta]), where the sum of these three cone populations at a given distance from the optic nerve (ON) head equals 100%. Black mouse: pigmented mouse strain, white mouse: albino mouse strain.

### 2.2. Despite the vast difference in S-opsin expression pattern, the distribution of true S-cones is strikingly similar between the pigmented and albino mouse

Such a highly skewed distribution of true S-cones conflicts with current understanding about the mouse retina. Therefore, we also examined an albino mouse line to determine whether this observation persists across different mouse strains. Interestingly, S-opsin expression extended well into the dorsal retina of the albino mouse (Fig. 1B-C) (Ortín-Martínez et al., 2014). Consequently, most cones in the dorsal retina were mixed cones, and true M-cones were very sparse (7%, compared to 97% in pigmented mouse, Fig. 1C right, Table S1A, Fig. 2A right). However, the percentage and distribution of true S-cones were remarkably conserved between strains (33% vs 29%, Fig. 1C and Table S1A) with nearly identical density maps (Fig. 2A, bottom row). The distribution of three main cone populations in four retinal quadrants centered upon the optic nerve head reveals different profiles for mixed and true M-cones, but a very similar pattern for true S-cones between the two mouse strains (Fig. 2C).

### 2.3. S-cone bipolar cells exhibit a dorsal-ventral gradient with a higher density in the ventral retina

One concern regarding cone classification based on opsin immunolabeling is that some S^+^M^-^ cones may instead be mixed cones with low M-opsin expression. In fact, a similar cone-type distribution of mouse retina has been observed; however, out of caution, S^+^M^-^ cones were only referred to as “anatomical” S-cones due to a lack of confirmation regarding their bipolar connections (Baden et al., 2013). Thus, both true S-cones and S-cone bipolar cells have been generally believed to be evenly distributed across the retina (Baden et al., 2013; Haverkamp et al., 2005; Wang et al., 2011). In order to confirm the distribution of true S-cones, it will be critical to uncover the distribution and dendritic contacts of S-cone bipolar cells (type 9 or SCBCs). Previously, SCBCs have only been identified among other bipolar, amacrine and ganglion cells in a Thy1-Clomeleon mouse line, rendering the quantification of their distribution across the entire retina impractical (Haverkamp et al., 2005). We generated a Copine9-Venus mouse line (Fig. 3, Table S1C), in which SCBCs are specifically marked, owing to the fact that *Cpne9* is a SCBC-enriched gene (Shekhar et al., 2016). These bipolar cells are often seen to extend long dendrites to reach true S-cones, bypassing other cone types (Fig. 3B-C). The majority of dendritic endings form enlarged terminals beneath true S-cones pedicles, but occasional slender “blind” endings are present (see arrows in Fig. 3C), which have been documented for S-cone bipolar cells in many species (Haverkamp et al., 2005; Herr et al., 2003; Kouyama and Marshak, 1992). Unexpectedly, we found that the distribution of SCBCs was also skewed toward VN retina, albeit with a shallower gradient (Fig. 3D-E). Thus, in the dorsal retina, the true S-cone to SCBC ratio is approximately 1:3.6, compared to 5.3:1 in the ventral retina (Table S2). Accordingly, we observed substantial divergence in the dorsal retina, with a single true S-cone connecting to as many as six SCBCs, whereas in the ventral retina, a single SCBC contacted 4-5 true S-cones (Fig. 3C). Thus, while SCBCs in VN retina are not as concentrated as true S-cones, they form convergent contacts exclusively with true S-cones. This confirms the identity of true S-cones revealed by immunohistochemistry and supports the finding that true S-cones are highly concentrated in VN mouse retina.

**Figure 3.**
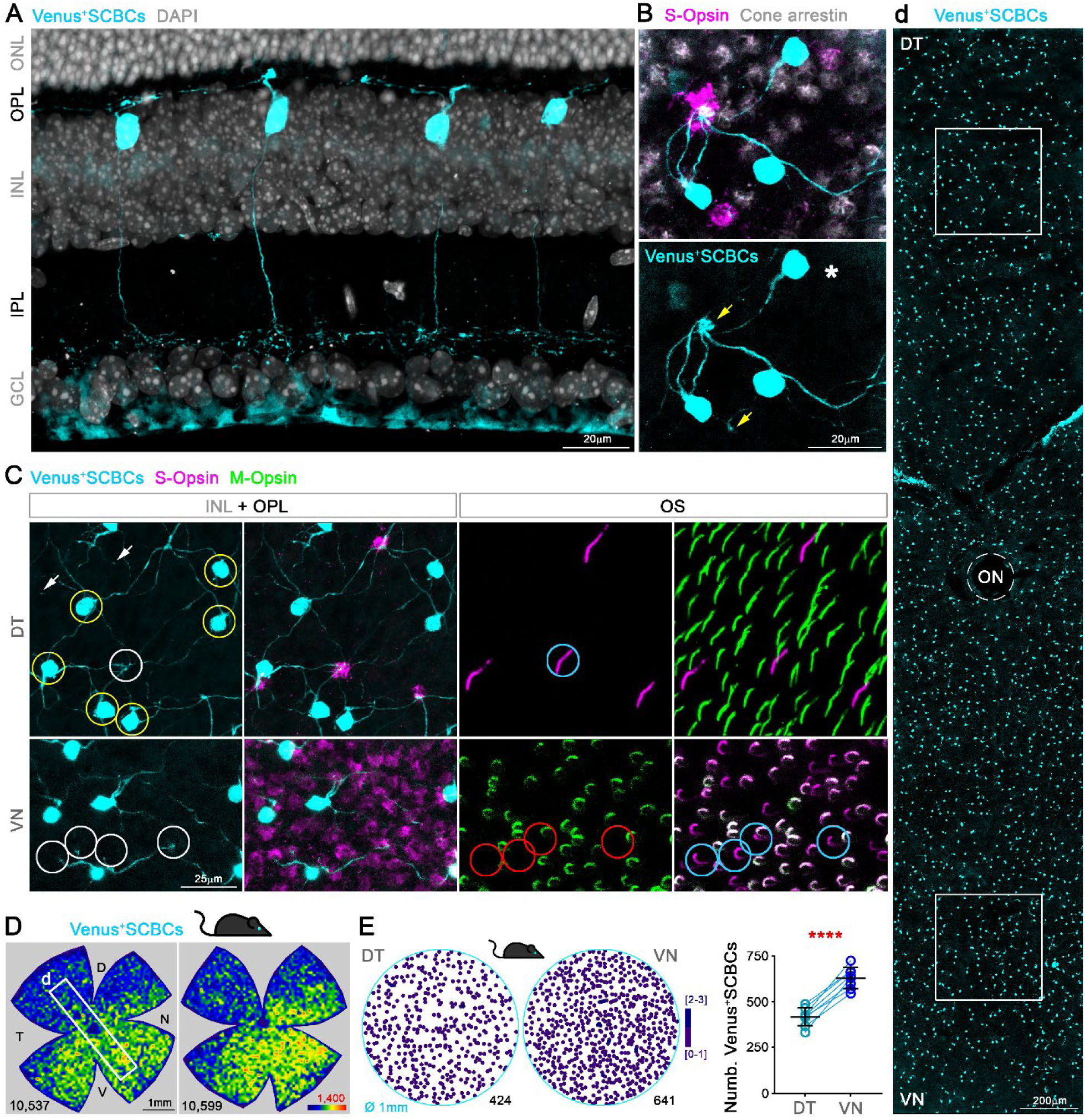
S-cone Bipolar cells (SCBCs) in Cpne9-Venus mouse retina. (A) Retinal cross section showing the characteristic morphology of SCBCs (Behrens et al., 2016; Breuninger et al., 2011). (B) Detailed view of the selective connectivity between Venus^+^SCBCs and true S-cone terminals (yellow arrows). Note that SCBCs avoid contacts with cone terminals lacking S-opsin expression (true M-cone pedicles, identified using cone arrestin), as well as a mixed cone pedicle, marked with an asterisk. (C) Images from flat-mount retinas focused on the inner nuclear and outer plexiform layers (INL+OPL) or in the photoreceptor outer segment (OS) layer of the corresponding area. Magnifications showing divergent and convergent connectivity patterns from true S-cone pedicles in dorsal and ventral retinal domains, respectively. In the DT retina, six Venus^+^SCBCs (yellow circles) contact a single true S-cone pedicle (white circle in DT); while one Venus^+^ SCBC contacts at least four true S-cone pedicles in the VN retina (white circles in VN), which belong to cones possessing S^+^M^-^OSs (blue circles). (D) Density maps depicting the distributions of SCBCs in Cpne9-Venus mice. (d) Venus^+^SCBCs along the DT-VN axis from a flat-mount retina (corresponding to the white frame in D) showing the gradual increase of SCBCs in VN retina where true S-cone density peaks (last row in Fig. 2A). (E) Demonstration of Venus^+^SCBC densities color-coded by the k-nearest neighbor algorithm according to the number of other Venus^+^SCBCs found within an 18 μm radius in two circular areas of interest (DT and VN). Venus^+^ SCBCs were significantly denser in VN (*p*<0.0001).

### 2.4. True S-cones in the ventral retina are not evenly distributed but form clusters

Given that the increased density of SCBCs in the ventral retina does not match that of true S-cones, individual SCBCs may be required to develop more dendrites to contact true S-cones. Intriguingly, we discovered in both strains that true S-cones in the ventral retina appeared to cluster together rather than forming an even distribution, as revealed by K-nearest neighbor analysis (Fig. 4A-B, Table S2). To further quantitatively assess how much they differ from a spatially even distribution, we computed two measures of regularity for true S cones: nearest neighbor and Voronoi diagram regularity indices (NNRI and VDRI, respectively (Reese and Keeley, 2015; Fig. 4C-D). Far from being evenly distributed, true S-cone placement was quite irregular and nearly indistinguishable from random placement (including a slight trend toward regularity measures lower than random, which may indicate a tendency toward clustering; see Reese, 2008). To further probe the possibility of true S-cone clustering, we investigated the proportions of true S-cone neighbors that are also true S-cones (denoted here as the S-cone neighbor ratio [SCNR]). Intriguingly, SCNRs were significantly larger than random chance—especially so in ventral retinas, further indicating a clustering of true S-cones in those areas (Fig. 4E). Notably, a more extreme form of clustering of S-cones has been observed in the “wild” mouse (Warwick et al., 2018) and with much lower densities in some felids (Ahnelt et al., 2000). Here, such clustering may reflect the mode of true S-cone development in the ventral retina, for example, by “clonal expansion” to achieve unusually high densities (Bruhn and Cepko, 1996; Reese et al., 1999). Intriguingly, it may also facilitate the wiring of true S-cones with sparsely distributed SCBCs in the ventral retina.

**Figure 4.**
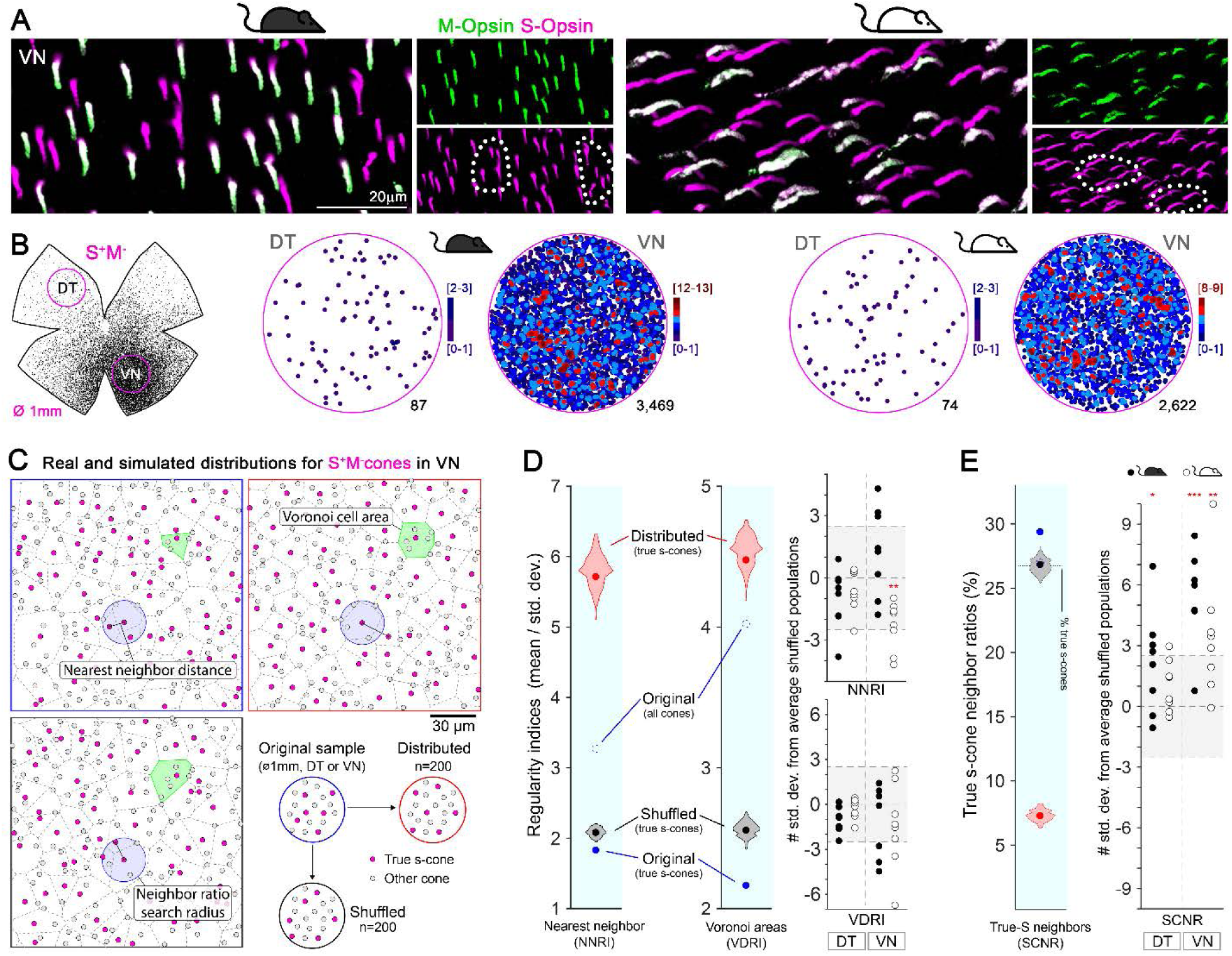
Clustering of true S-cones in the ventronasal (VN) retina. (A) Retinal magnifications from flat-mount retinas demonstrating grouping of true S-cones in the VN area, where true S-cone density peaks. Dashed lines depict independent groups of true S-cones that are not commingled with mixed cones (M^+^S^+^, white outer segments in the merged image). (B) Retinal scheme of true S-cones used for selecting two circular areas of interest along the dorsotemporal-ventronasal (DT-VN) axis. Circular maps demonstrate true S-cone clustering in these regions. true S-cone locations are color-coded by the k-nearest neighbor algorithm according to the number of other true S-cones found within an 18 μm radius. (C-E) Analytical comparisons of DT and VN populations of true S-cones to their simulated alternatives. C) Example real and simulated true S-cone populations and their quantification. Images depict true S-cone locations (magenta dots) and boundaries of their Voronoi cells (dashed lines) from original and example simulated (“distributed”, “shuffled”) cone populations. Gray dots indicate the locations of other cone types. Observed cone locations were used for all simulated populations; only their cone identities were changed. The annotated features are examples of those measurements used in the calculations presented in D-E. (D) Comparison of sample regularity indices for one albino VN retinal sample to violin plots of those values observed for n=200 simulated cone populations. Note that average regularity indices for true S-cones were lower than that of shuffled populations, whereas those values lay between shuffled and distributed populations when all cones were considered. Plots on the right show values for all actual retinal samples normalized using the mean and standard deviations of their simulated “shuffled” counterparts. The y-axis range corresponding to ± 2.5 standard deviations from the mean (i.e., that containing ∼99% of shuffled samples) is highlighted. (E) Comparison of the real average SCNR for the example in C-D to those values for its simulated counterparts. Note that the average SCNR for all cones in this sample was equal to that predicted by random chance (i.e., the ratio of true S-cones to all cones), which in turn was equal to the average for true S-cones for shuffled samples. In contrast, the real true S-cone SCNR was higher. Plot on the right shows true S-cone SCNR values for all samples, normalized as described for D.

### 2.5. Enriched true S-cones in the ventral retina may provide an anatomical base for mouse color vision

Despite having a rod-dominated retina, mice can perceive color (Denman et al., 2018; Jacobs et al., 2004). Although it remains uncertain whether the source of long-wavelength sensitive signals for color opponency arises in rods or M-cones (Baden and Osorio, 2019; Ekesten et al., 2000; Ekesten and Gouras, 2005; Joesch and Meister, 2016; Reitner et al., 1991), it is clear that true S-cones provide short-wavelength signals for color discrimination. Thus, such high enrichment of true S-cones in the ventral retina is a previously missed anatomical feature for mouse color vision. From projections mapping true S-cone densities into visual space (Fig S2; Sterratt et al., 2013), it is conceivable that high ventral true S-cone density will provide a much higher sensitivity of short-wavelength signals, thus facilitating color detection for the upper visual field. Although the true S-cone signals carried by SCBCs in the dorsal retina might not be significant for color detection, they could certainly participate in other functions, such as non-image forming vision, that are known to involve short-wavelength signals (Altimus et al., 2008; Doyle et al., 2008; Patterson et al., 2020). Interestingly, the overall true S-cone percentage in the mouse retina remains approximately 10% (Fig. 2B), and the average true S-cone to SCBC ratio across the whole retina is about 1.7:1 (Table. 1B-C), similar to what has been reported in other mammals (Ahnelt et al., 2006; Ahnelt and Kolb, 2000; Hendrickson and Hicks, 2002; Hunt and Peichl, 2014; Kryger et al., 1998; Lukáts et al., 2005; Müller and Peichl, 1989; Ortín-Martínez et al., 2010; Peichl et al., 2000; Schiviz et al., 2008; Shinozaki et al., 2010). Such a spatial rearrangement of true S-cones and SCBCs likely reflects evolutionary adaption to enhance color coding for the upper visual field as best suited for mice’s habitat and behavior. In addition, the clustering of true S-cones in the ventral retina may allow several neighboring cones to converge onto the same BC, thus enhancing signal-to-noise ratios for more accurate detection (Schmidt et al., 2019). It is also remarkable that despite the very different S-opsin expression patterns, the true S-cone population and distribution are strikingly similar between pigmented and albino mice, suggesting a common functional significance.

## 3. ACKNOWLEDGEMENTS

The authors would like to thank the NEI Animal Care team, especially Megan Kopera and Ashley Yedlicka.

## 4. AUTHOR CONTRIBUTIONS

The experiments were conceived and designed by F.M.N.N., V.P.K. and W.L. Experiments were conducted by F.M.N.N., B.P., A.K., G.Z. and data were analyzed by F.M.N.N., B.P., A.K., J.B., G.Z., W.L. The Copine9-Venus mouse line was generated by L.D. The automatic script and simulations were written by F.M.N.N. and J.B. The original draft was written by F.M.N.N. and W.L. and all authors have written, reviewed and edited the final paper.

## 8. SUPPLEMENTARY MATERIAL

**Table S1.**
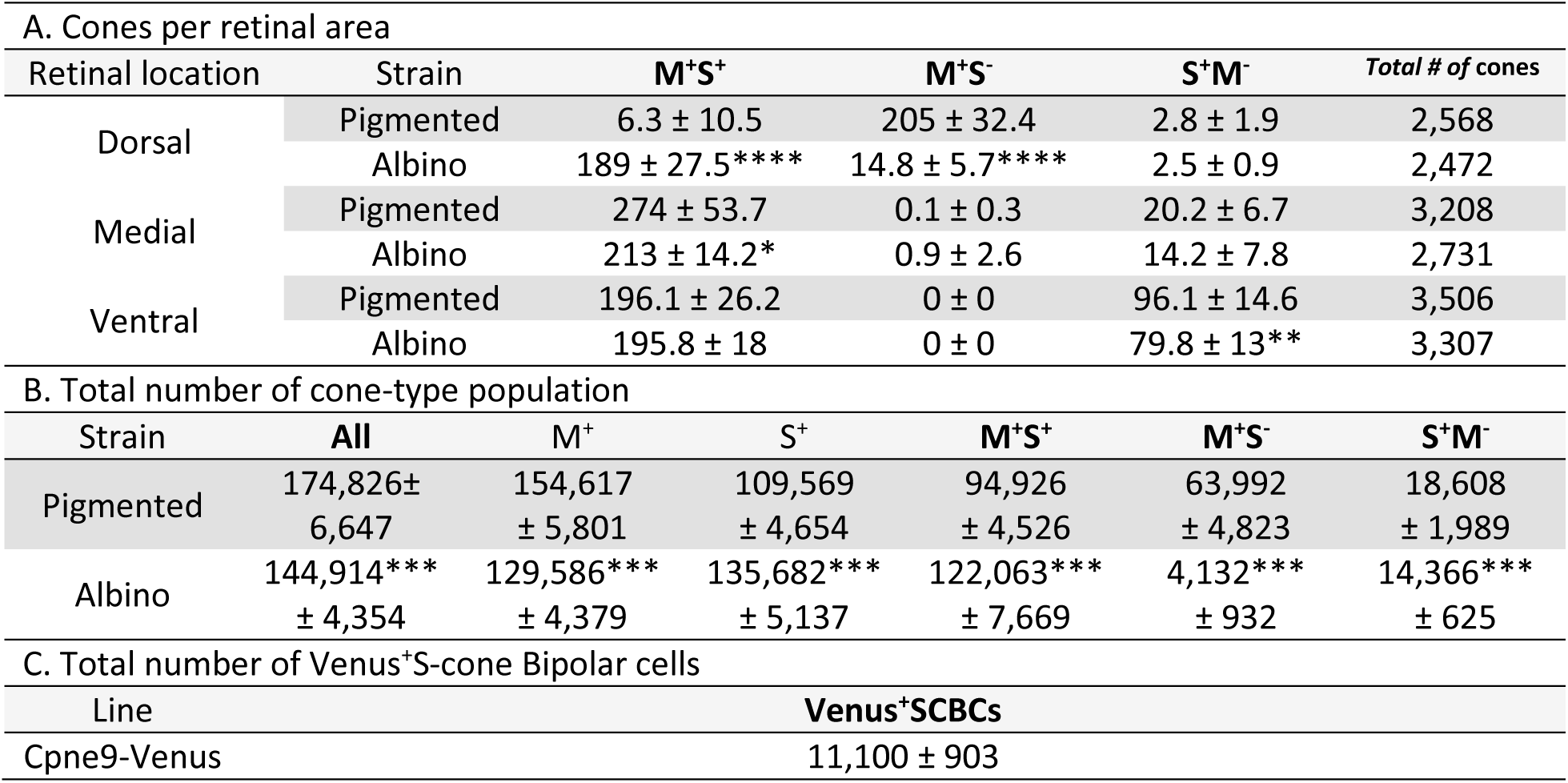
(A) Cone numbers in different retinal areas along the dorsoventral axis in pigmented and albino mouse. Three images/area (dorsal, medial and ventral) from four retinas/strain. Different cone type quantifications are shown as average ± SD, corresponding to the percentages shown in Fig 1C. The total number of cones analyzed per location and strain are shown in the last column. Total number of cones (B) or S-cone Bipolar cells (SCBCs, C) in eight retinas/mouse strain or line (average ± SD, see also Fig. 2B). Significant differences between strains *p*<0.05 (*), *p*<0.01 (**), *p*<0.001 (***), *p*<0.0001 (****).

**Table S2.** Numbers of true S-cones (A) and Cpne9-Venus^+^SCBCs (B) in dorsotemporal (DT) and ventronasal (VN) circular areas (1mm diameter, Figs 3E and 4B). Quantitative data are shown as average ± SD from eight retinas/strain or line. The mean of true S-cones and Venus^+^SCBCs in these circular areas was used to calculate the DT:VN and true S-cone:SCBC (C) ratios. Significant differences between strains *p*<0.05 (*), *p*<0.001 (***). True S-cones and SCBCs were significant different between DT and VN retina (*p*<0.0001).

| Strain/Line | DT | VN | DT:VN Ratio |
| --- | --- | --- | --- |
| A. true S-cones |  |  |  |
| Pigmented | 115 ± 34 | 3,359 ± 437 | 1:31 |
| Albino | 82 ± 10* | 2,346 ± 405*** | 1:29 |
| B. S-Cone Bipolar Cells |  |  |  |
| Cpne9-Venus | 417 ± 49 | 630 ± 58 | 1:1.5 |
| C. true S-cone:S-cone bipolar cell Ratio |  |  |  |
| Pigmented | 1:3.6 | 5.3:1 |  |
| Albino | 1:5.1 | 3.7:1 |  |

**Figure S1.**
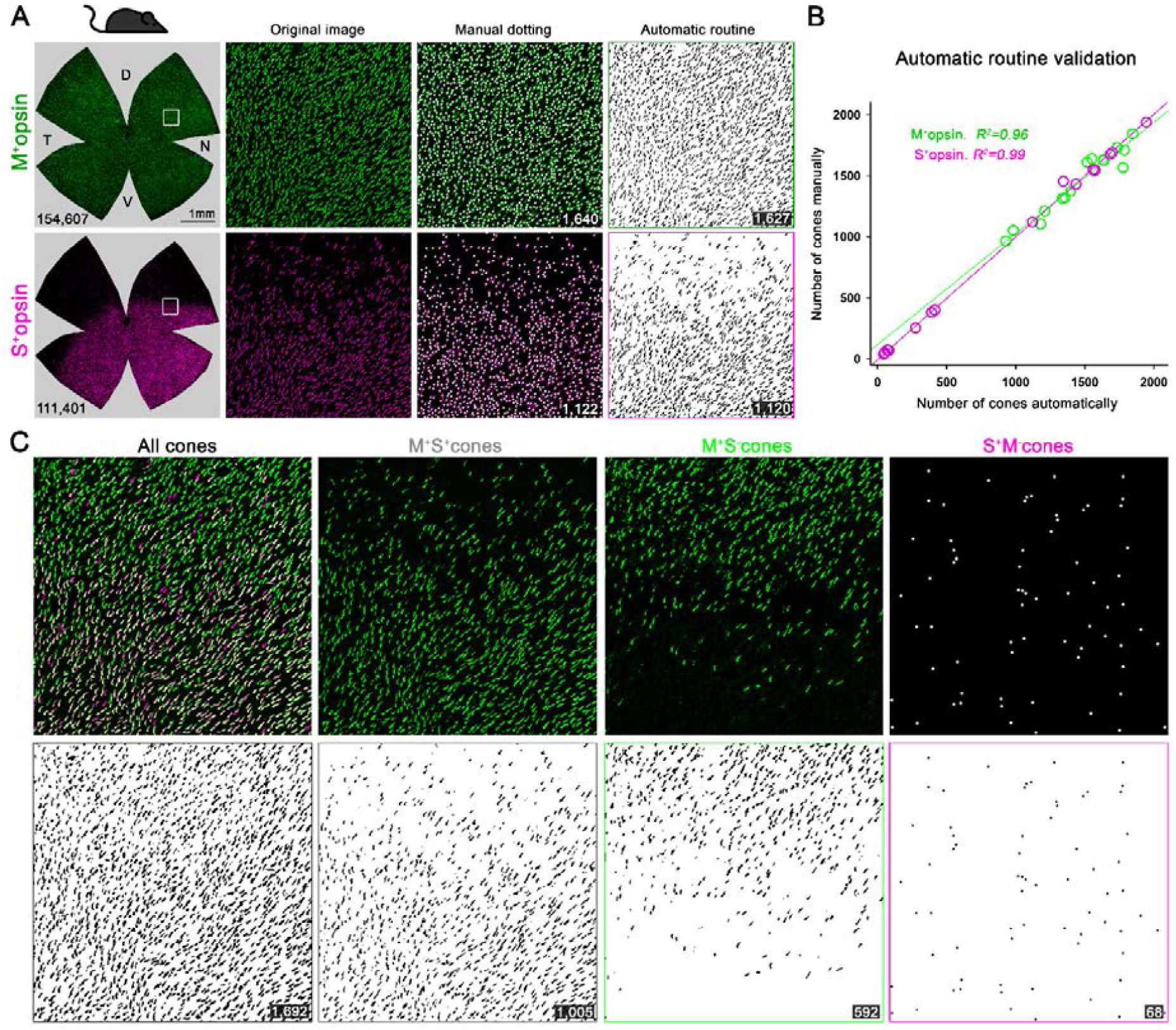
Validation of automatic routine for cone outer segment quantification. (A) Retinal photomontages for M- and S-opsin signal in the same pigmented retina (correspond to second column in Fig. 2A). The square depicts an area of interest selected (transition zone of S-opsin expression) to perform the automatic routine validation by comparing manual and automatic quantifications. The images processed by the automatic routine using ImageJ show the selection of positive objects from the corresponding original image. (B) X, Y graph showing the linear correlation (Pearson coefficient, *R*^*2*^) between manual and automatic quantifications. 21,898 M^+^ and 13,705 S^+^cones were manually annotated while 21,689 M^+^ and 13,661 S^+^cones were automatically identified in 3 random images obtained from 5 retinal photomontages. (C) All, mixed, true M- and true S-cone populations are extracted from the original M- and S-cone images. All-cones were quantified after overlapping M- and S-signals. mixed (M^+^S^+^) cones were obtained by subtracting the background of the S-opsin image in the M-opsin one. true M-cones (M^+^S^-^) for pigmented mice are obtained after subtracting the S-opsin signal to the M-opsin photomontage. Finally, true M-cones for albino and true S-cones (S^+^M^-^), in both strains, are manually marked on the retinal photomontage (Adobe Photoshop CC). The B&W images shown the processed image after quantifying automatically. At the bottom of each image is shown the number of quantified cones. Black mouse: pigmented mouse strain.

**Figure S2.**
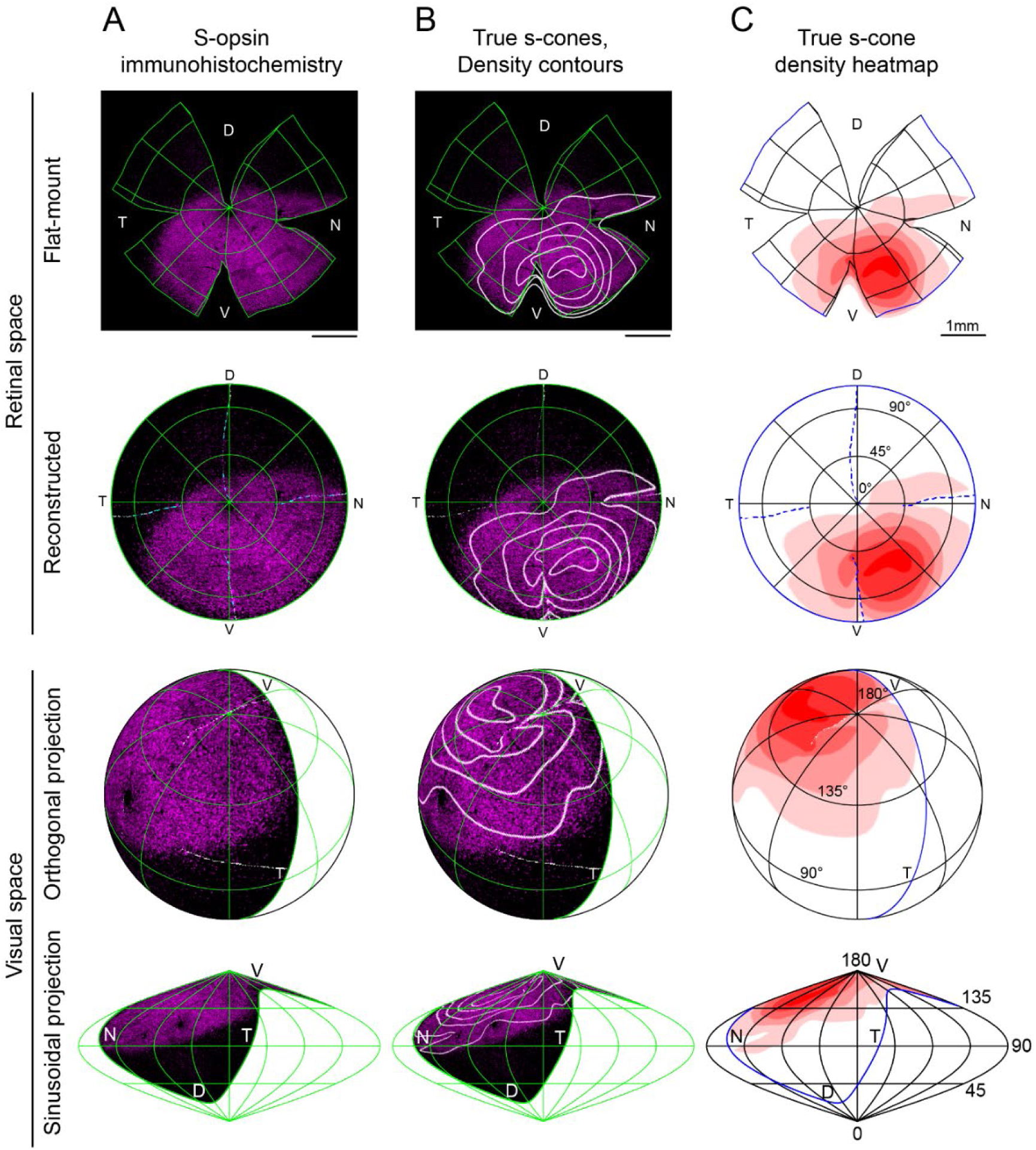
Reconstruction and mapping of true S-cone densities into visual space. Representative left eye from a 3-month-old pigmented mouse (C57). (A) S-opsin antibody labeling; (B) true s-cone density contour lines separated by quintiles overlaid onto s-opsin labeling; (C) quintile heatmap contours of true s-cone density. The top two rows demonstrate the flat-mount retina with marks for edges and relaxing cuts, followed by its reconstruction into uncut retinal space with lines of latitude and longitude that have been projected onto the flat-mount. The bottom two rows show the reconstructed retina inverted into visual space using orthogonal and sinusoidal projections. For these views, eye orientation angles for elevation and azimuth of 22° and 64°, respectively, have been used as in (Sterratt et al., 2013). For orthogonal projections, the globe has been rotated forward by 50° to emphasize the relationship of true S-cone densities to the upper pole of the visual field. S-opsin labeling is restricted to the upper visual field, but true S-cones are concentrated toward its lateral edges.

## 10. METHODS

### 10.1. KEY RESOURCES

| REAGENT or RESOURCE | SOURCE | IDENTIFIER |
| --- | --- | --- |
| **Antibodies** |  |  |
| Goat polyclonal anti-OPN1SW (N-20) | Santa Cruz Biotechnology | Cat#sc-14363 |
| Rabbit polyclonal anti-Opsin Red/Green | Millipore/Sigma | Cat#AB5405 |
| Rabbit polyclonal anti-Cone arrestin | Millipore/Sigma | Cat#AB15282 |
| Chicken polyclonal anti-GFP | Millipore/Sigma | Cat#AB16901 |
| Donkey polyclonal anti-Rabbit 488 | Jackson ImmunoResearch | Cat#711-547-003 |
| Donkey polyclonal anti-Rabbit Cy3 | Jackson ImmunoResearch | Cat#711-165-152 |
| Donkey polyclonal anti-Goat 647 | Jackson ImmunoResearch | Cat#705-605-147 |
| Donkey polyclonal anti-Goat Cy3 | Jackson ImmunoResearch | Cat#705-166-147 |
| Donkey polyclonal anti-Chicken 488 | Jackson ImmunoResearch | Cat#703-545-155 |
| **Experimental Models: Organisms/Strains** |  |  |
| C57BL/6J pigmented mouse inbred strain | Jackson Laboratory | Cat#000664 |
| CD-1 IGS albino mouse strain | Charles River | Cat#022 |
| Copine9-Venus mouse line | This paper | - |
| **Software and Algorithms** |  |  |
| Zen lite Black edition 2.3 SP1 | Zeiss | https://www.zeiss.com/microscopy/us/products/microscope-software/zen-lite.html |
| FIJI-ImageJ v1.52r | NIH | https://imagej.nih.gov/ij/ |
| Sigma Plot 13.0 | Systat Software | https://systatsoftware.com/products/sigmaplot/ |
| GraphPad Prism 8.3.0 | Graph Pad Software | https://www.graphpad.com/scientific-software/prism/ |
| Photoshop CC 20.0.6 | Adobe | https://www.adobe.com/products/photoshop.html |
| MATLAB2016 | MathWorks | https://www.mathworks.com/products/matlab.html |
| R | The R Project for Statistical Computing | https://www.r-project.org/ |
| Retina and Visual Space Retistruct Package | Sterratt DC et al., PLoS Comput Biol 9(2): e1002921 | https://journals.plos.org/ploscompbiol/article?id=10.1371/journal.pcbi.1002921 |
| Zotero 5.0 | Corporation for Digital Scholarship | https://www.zotero.org/download/ |

### 10.2. LEAD CONTACT AND MATERIALS AVAILABILITY

Further information and requests for resources and reagents should be directed to and will be fulfilled by the Lead Contact, Wei Li.

### 10.3. METHOD DETAILS

#### 10.3.1. Animal handling and Ethic statement

Three months old male pigmented (*C57BL/6J*, n=5), albino (*CD1*, n=5) mice were obtained from the National Eye Institute breeding colony. The Venus-Cpne9 mouse line (based on previous single cell sequencing data (Shekhar et al., 2016)) carries a reporter (Venus) allele under the control of the mouse Cpne9 locus. The reporter allele was created directly in B6.SJL(F1) zygotes using CRISPR-mediated homologous recombination (HR) (Yang et al., 2013). Briefly, a HR targeting template was assembled with PCR fragments of 5’ and 3’ homology arms of 910 bp and 969 bp respectively, flanking exon one, and a Venus expression cassette carrying the bovine growth hormone polyadenylation (bGH-PolyA) signal sequence as the terminator. Homology arms were designed such that integration of the reporter cassette would be at the position right after the first codon of the Cpne9 gene in exon one. A pair of guide RNAs (gRNA), with outward orientation (38 bp apart), were synthesized by *in vitro* transcription as described (Yang et al., 2013) and tested for their efficiency and potential toxicity in a zygote differentiation assay where mouse fertilized eggs were electroporated with SpCas9 protein and gRNA ribonuclear particles. Eggs were cultured in vitro for 4 days in KSOM (Origio Inc, CT) until differentiated to blastocysts. Viability and indel formation were counted respectively. gRNA sequences are (1) Copine9_gRNA_L(73/25), 5’GAGACATGACTGGTCCAA3’; (2) Copine9_gRNA_R(62/4.40), 5’GCCTCGGAGCGTAGCGTCC3’. A mixture of the targeting plasmid (super coiled, 25ng/µl) with two tested gRNAs (25 ng/µl each) and the SpCas9 protein (Life Science technology, 30ng/µl) were microinjected into mouse fertilized eggs and transferred to pseudopregnant female recipients as described elsewhere (Yang et al., 2013). With a total of 15 F0 live births from 6 pseudopregnant females, 11 were found to carry the knockin allele by homologous recombination, a HR rate of 73%. F0 founders in B6.SJL F1 (50% C57BL6 genome) were crossed consecutively for 3 generations with C57BL6/J mice to reach near congenic state to C57BL6/J.

Mice were housed a 12:12 hours light/dark cycle. All experiments and animal care are conducted in accordance with protocols approved by the Animal Care and Use Committee of the National Institutes of Health and following the Association for Research in Vision and Ophthalmology guidelines for the use of animals in research.

#### 10.3.2. Tissue collection

All animals were sacrificed with an overdose of CO2 and perfused transcardially with saline followed by 4% paraformaldehyde. To preserve retinal orientation, eight retinas per mouse strain were dissected as flat whole-mounts by making four radial cuts (the deepest one in the dorsal pole previously marked with a burn signal as described (Nadal-Nicolás et al., 2018; Stabio et al., 2018). The two remaining retinas were cut in dorso-ventral orientation (14μm) after cryoprotection in increasing gradients of sucrose (Sigma-Aldrich SL) and embedding in optimal cutting temperature (OCT; Sakura Finetek).

#### 10.3.3. Immunohistochemical labeling

Immunodetection of flat-mounted retinas or retinal sections was carried out as previously described (Nadal-Nicolás et al., 2018). Importantly, the retinal pigmented epithelium was removed before the immunodetection. First, whole-retinas were permeated (4×10’) in PBS 0.5% Triton X-100 (Tx) and incubated by shaking overnight at room temperature with S-opsin (1:1200) and M-opsin (1:1000) or cone arrestin (1:300) primary antibodies diluted in blocking buffer (2% normal donkey serum). Cpne9-Venus retinas were additionally incubated with an anti-GFP antibody (1:100) to enhance the original Venus signal. Retinas were washed in PBS 0.5% Tx before incubating the appropriate secondary antibodies overnight (1:500). Finally, retinas were thoroughly washed prior to mounting with photoreceptor side up on slides and covered with anti-fading solution. Retinal sections were counterstained with DAPI.

#### 10.3.4. Image acquisition

Retinal whole-mounts were imaged with a 20x objective using a LSM 780 Zeiss confocal microscope equipped with computer-driven motorized stage controlled by Zen Lite software (Black edition, Zeiss). M- and S-opsins were imaged together to allow the identification and quantification of different cone types. Magnifications from flat mounts and retinal cross-sections (Fig. 1) were taken from dorsal, medial and ventral areas using a 63x objective for opsin co-expression analysis. Images from retinal cross-sections were acquired ∼1.5mm dorsally or ventrally from the optic disc.

#### 10.3.5. Sampling and opsin co-expression measurement

In four retinas per strain, we acquired images from three 135×135 μm samples (63x) per each area of interest (dorsal, medial and ventral). These areas were selected according to the S-opsin gradient in wholemount retinas (see scheme in Fig. 1C). Cone outer segments were manually classified as true M- (M^+^S^-^), true S- (S^+^M^-^) or mixed (M^+^S^+^) cones depending on their opsin expression. Data representation was performed using GraphPad Prism 8.3 software.

#### 10.3.6. Image processing: manual and automated whole quantification

To characterize the distribution of the different cone photoreceptor types in the mouse retina, we developed and validated an automatic routine (ImageJ, NIH) to identify, quantify the total number of outer segments and finally extract the location of each individual cone (Fig. S1A).

Briefly, maximum-projection images were background-subtracted and thresholded to create a binary mask that was then processed using watershed and despeckle filters to isolate individual cones and reduce noise. The “3D Objects Counter” plugin was applied to such images to count cones within fixed parameters (shape and size) and extract their *xy* coordinates for further analysis. This automation was validated by statistical comparation with manual counting performed by an experienced investigator (Pearson correlation coefficient *R*^2^ = 96-99% for M- or S-opsin respectively, Fig. S1B). To count cone subtypes, images were pre-processed with image processing software (Adobe Photoshop CC) to isolate the desired subtype and then manually marked using Photoshop, or automatically counted using ImageJ as described above. Total cone populations were determined by combining M- and S-opsin channels, while mixed M^+^S^+^ cones were obtained by masking the M-opsin signal with the S-opsin channel. true M-cones in pigmented mice were obtained by subtracting the S-opsin signal from the M-opsin photomontage. Finally, true M-cones (in albino samples), true S-cones (both strains) (Fig. S1C) and Venus^+^SCBCs (Cpne9-Venus mouse line) were manually marked on the retinal photomontage (Adobe Photoshop CC).

#### 10.3.7. Topographical distributions

Topographical distributions of cone population densities were calculated from cone locations identified in whole-mount retinas using image processing (see above). From these populations, isodensity maps were created using Sigmaplot 13.0 (Systat Software). These maps are filled contour plots generated by assigning to each area of interest (83.3×83.3 μm) a color code according to its cone density, ranging from 0 (purple) to 17,300 cones/mm^2^ for all cone types except for true S-cones and true M-cone in the albino strain (5,000 cones/mm^2^), as represented in the last image of each row of Fig. 2A, or 1,400 SCBCs/mm^2^ (Fig. 3D) within a 10-step color-scale. These calculations allow as well, the illustration of the number of cones at a given position from the ON center. To analyze the relative opsin expression along the retinal surface, we have considered three cone populations (mixed, true M- and true S-cones) dividing the retina in four quadrants: dorsotemporal, dorsonasal, ventrotemporal and ventronasal (DT, DN, VT and VN respectively, scheme in Fig. 2C). The relative percentage of cone-types are represented in line graphs from four retinas/strain (SigmaPlot 13.0).

#### 10.3.8. SCBC sampling and ‘true S-cone’ connectivity

To characterize the connectivity of Venus^+^S-cone bipolar cells (Venus^+^SCBCs) with true S-cone terminals, we acquired images from the same area (260×260 μm, 63x) at two focal planes: First, we focused upon the INL+OPL, then the corresponding photoreceptor outer segment (OS) layer, respectively, for two areas of interest (DT and VN). To verify connectivity between Venus^+^SCBCs dendrites and true S-cone pedicles in the OPL, we also stained the cone pedicles with cone arrestin and traced cone pedicles to their respective OS to verify S^+^M^-^ opsin labeling (Fig. 3).

#### 10.3.9. Clustering analysis. K-neighbor maps and variance analysis of Voronoi dispersion

To assess the clustering of true S-cones and S-cone bipolar cell (SCBC) clustering, we performed two comparable sets of analyses. First, we extracted two circular areas (1mm diameter) in the DT-VN axis at 1mm from the optic disc center (scheme in 4B). A K-nearest neighbor algorithm (Nadal-Nicolás et al., 2014) was used to map the number of neighboring true S-cones within a 18 μm radius of each true S-cone to a color-code in its retinal position (Fig. 4B). Regularity indices were computed for each retinal sample using Voronoi diagrams for cone positions as well as nearest neighbor distances (VDRI and NNRI, respectively (Reese and Keeley, 2015); Fig. 4C-E). NNRIs were computed as the ratio of the mean to the standard deviation for the distance from true S-cones to their nearest true S-cone neighbor. true S-cone neighbor ratios (SCNR) were calculated for each retinal sample as the average proportion of true s-cones within a given radius for each cone. This search radius was calculated separately for each sample to correct for sample-to-sample variations in total density: this radius (*r*) was calculated as *r* = 3√ (*A* / (√2 *πN*)), where *A* is the circular area of the 1mm diameter retinal sample and *N* is the total number of cones in that sample. For a highly regular cell mosaic containing *N* cells filling an area *A*, this calculation estimates the location of the first minimum in the density recovery profile (Rodieck, 1991), providing the average radius of a circle centered upon a cone that will encompass its first tier of cone neighbors (but exclude the second tier) in an evenly distributed mosaic. To minimize edge effects from computations of NNRI, VDRI, SCNR, those values for cones closer to the outer sample edge than the SCNR search radius were discarded. To produce simulated cone mosaics for comparison with observed values, cone distributions with evenly “distributed” true S-cones were generated by first using a simple mutual repulsion simulation to maximize the distances between true S-cones, followed by assigning the nearest positions among all cone locations as being “true S”. “Shuffled” populations of true S-cones were generated by permuting cone identities randomly among all cone locations, holding the proportion of true S-cones constant. Voronoi diagrams, neighbor calculations, and mosaic generation and other computations were performed using MATLAB R2016b.

#### 10.3.10. Retinal reconstruction and visuotopic projection

Retinal images were reconstructed and projected into visual space using R software v.3.5.2 for 64-bit Microsoft Windows using Retistruct v.0.6.2 as in (Sterratt et al., 2013). Reconstruction parameters from that citation were used: namely, a rim angle of 112° (phi0 = 22°), and eye orientation angles of 22° (elevation) and 64° (azimuthal). For Figure S2, true s-cone density contour lines and heatmaps were computed in MATLAB and overlaid onto flat-mount retina opsin labeling images using ImageJ prior to processing by Retistruct.

#### 10.3.11. Statistical analysis

Statistical comparisons for the percentage of cones/retinal location, the total cone quantifications (Table S1) and the DT or VN true S-cones and Venus^+^SCBCs (Table S2) were carried out using GraphPad Prism v8.3 for Microsoft Windows. Data are presented as mean ± standard deviation. All data sets passed the D’Agostino-Pearson test for normality, and the comparisons between strains were performed with Student’s *t*-test.

For each 1mm retinal sample, VDRI, NNRI, and SCNR values were normalized and compared to the distributions of “shuffled” cone populations. Such comparisons were not performed against “distributed” populations, because in those populations, VDRI and NNRI values were consistently much higher—and SCNR much lower—than in real samples (see Fig. 4D-E). The “shuffled” populations for each retinal region produced measurements that were well described by normal distributions (Kolmogorov-Smirnov test, MATLAB). Thus, to allow comparisons across samples, we converted each measurement into a *Z*-score using the mean and standard deviation of those measures from shuffled populations. One-tailed Student’s *t*-tests were performed to compare the normalized measures to the distribution of “randomly shuffled” cone population measures, and significance was determined at the *p*<0.05 level.

